# Therapeutic window for treatment of prion disease defined by timed depletion of Prion protein

**DOI:** 10.1101/2024.11.18.624122

**Authors:** Barry M. Bradford, Alison Marshall, Nadia L. Tuzi, Deborah Brown, Dorothy Kisielewski, Kim M. Summers, V. Hugh Perry, Pedro Piccardo, Alan R. Clarke, Jean C. Manson

## Abstract

There is no effective treatment preventing the progression of neurodegenerative diseases such as prion and Alzheimer’s diseases. These fatal diseases of the central nervous system, involve progressive accumulation of a misfolded protein long before overt clinical signs of disease. Removal of prion protein early in the pathological process appears to halt the progression however, it is not known whether intervention at later disease stages could be effective. We investigated the potential for intervention throughout the course of prion disease, by developing a mouse model in which *Prnp* expression can be manipulated in a tissue specific and time dependent manner. Depleting *Prnp* from neuronal populations in CNS throughout the preclinical phase substantially prolonged incubation. The pathology was dramatically altered to a pattern of astrocytic associated prion deposition. However once overt clinical symptoms of disease were apparent *Prnp* depletion did not alter disease progression. This study establishes a wide window for intervention, and suggests timely treatment could delay the onset of clinical disease potentially well beyond the lifetime of an individual.

## Introduction

In prion disease the native cellular isoform of the prion protein (PrP^C^; uniprot P04925, encoded by the *Prnp* gene) is misfolded into the disease-associated form PrP^Sc^ (Prusiner, 1982). Clinical and pathological signs of disease are completely dependent upon the expression of PrP^C^. When wild type animals carrying the normal *Prnp* gene are infected with prion agents, they show accumulation and deposition of the protease resistant PrP^Sc^ in the brain, which can be detected very early in the disease process (Bencsik et al., 2008, Bruce et al., 1994, Schulz-Schaeffer et al., 2000). This is accompanied by characteristic deterioration in neuronal dendritic morphology including varicosities and a marked loss of spines (Brown et al., 2001, Jeffrey et al., 2000, Johnston et al., 1997). Synaptic degeneration occurs and eventually progresses to neuronal death with loss of neuronal soma (Belichenko et al., 2000). The central nervous system (CNS) grey matter develops a vacuolar spongiform appearance, hence the name transmissible spongiform encephalopathy (TSE). In contrast, transgenic animals lacking PrP^C^ expression due to *Prnp* gene knockout are wholly resistant to disease and show none of this neuropathology (Bueler et al., 1993, Manson et al., 1994a, Richt et al., 2007). Disease incubation period is modulated by the total amount of PrP^C^ in the brain, with PrP hemizygous mice showing an extended incubation period (Manson et al., 1994b, Scott et al., 1989). Moreover when neuronal PrP^C^ expression was depleted eight weeks post infection the disease process was arrested (Mallucci et al., 2003) with the reversal of both the characteristic spongiform pathology and behavioral deficits (Mallucci et al., 2007). Since the expression of PrP^C^ has such a dramatic influence on the disease process, manipulation of *Prnp* expression provides an ideal system to investigate stages at which that process can be moderated and hence determine the window in which therapy could modify disease progression.

We have altered the expression of the *Prnp* gene within the CNS to investigate the potential for disease intervention throughout the extended preclinical and clinical phase of disease. We developed a mouse model in which *Prnp* expression in neurons can be manipulated, by combining gene-targeted *LoxP* flanked *Prnp* controlled by the endogenous murine promoter (Tuzi et al., 2004) and tamoxifen-inducible neuronal expressed Cre recombinase transgenes to allow timed control of reduction of PrP^C^ from the CNS. We infected this model with the well-characterized ME7 and 139A mouse-adapted scrapie and 301C mouse-adapted bovine spongiform encephalopathy (BSE) strains of agent and showed that when CNS PrP^C^ was reduced prior to the clinical phase of infection, incubation times were extended. Neuropathological outcome was also strikingly altered, with preservation of neuronal cell bodies despite extensive astrocytic associated deposition of PrP^Sc^. We demonstrated a substantial prolongation of incubation time throughout the preclinical phase of disease, only becoming negligible when overt clinical symptoms of disease were apparent. Hence we have established a wide therapeutic window for intervention which could delay the clinical disease onset potentially well beyond the lifetime of an individual.

## Results

We have produced a transgenic mouse model expressing a tamoxifen-inducible Cre fusion protein (CreERT) expressed via the *Rattus norvegicus* neuron specific gamma enolase (*Eno2* gene; uniprot P07323) promoter. We confirmed that Cre activity in these NSECreERT mice was present only after tamoxifen induction using the ROSA26-LacZ reporter for Cre-mediated recombination. Beta-galactosidase activity was observed in distinct large bodied neurons of the central nervous system including hippocampal pyramidal cells (Fig. 1A), magnocellular neurons of the red nucleus (Fig. 1B) and cerebellar Purkinje cells (Fig. 1C). Extensive reporter activity was present throughout small soma neurons in the CNS including for example cerebellar granule cells (Fig. 1C & 1D) and small and large soma neurons in the brain stem (Fig. 1E) and grey matter of the spinal cord (Fig. 1F). The extent and intensity of beta-galactosidase activity was variable between the tamoxifen treated animals but no expression was detectable in animals which were not treated with tamoxifen.

**Figure 1.**
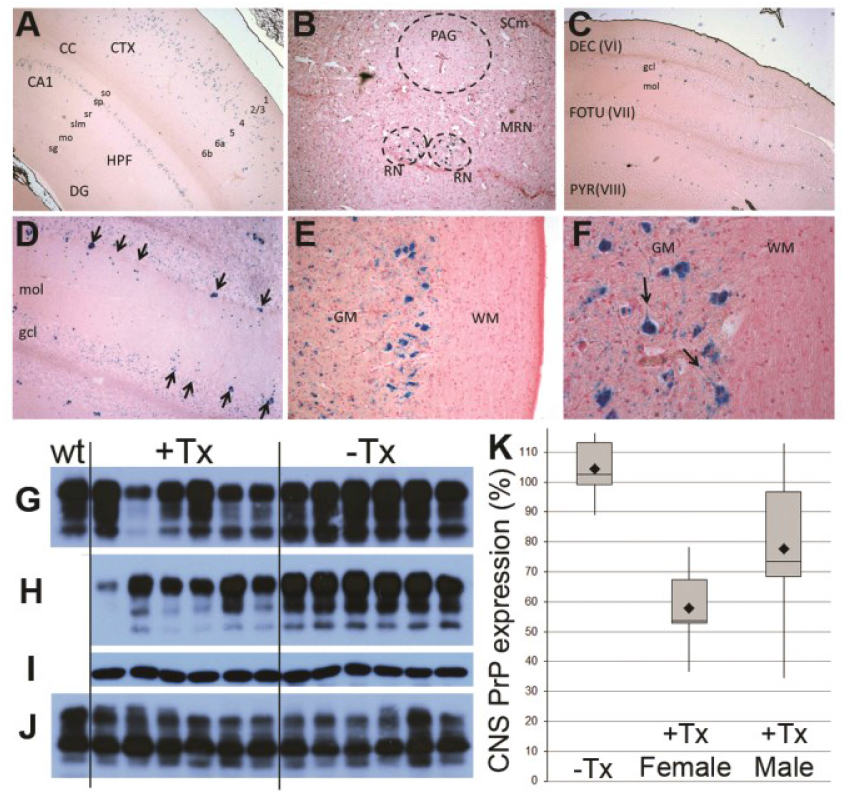
Characterization of tamoxifen-inducible Cre activity and neuronal PrP^C^ depletion using NSECreERT transgenic mice. Tamoxifen-inducible Cre activity resulted in β-galactosidase expression in ROSA26-LacZ reporter (A-F), detected by X-gal histochemistry (blue). (A) LacZ positive layered cortical neurons (CTX), and hippocampal formation (HPF) pyramidal neurons in the stratum pyrimidale (sp) original magnification x40. (B) LacZ positive magnocellular neurons of the red nucleus (RN) and widespread neurones in other midbrain regions MRN and SCm, little staining was observed in the periaqueductal grey region (PAG). (C) LacZ positive cerebellar Purkinje neurons, original magnification x40, (D) neurons in the granule cell layer (gcl) and cerebellar Purkinje neurones (arrows) show X-gal staining, the amount of Purkinje cell somatic and dendritic staining in the molecular layer (mol) varies from cell to cell, original magnification x100. (E) Neurons in the thoracic spinal cord grey matter (GM), no staining was observed in white matter (WM), original magnification x40. (F) Cytoplasmic localization of X-gal staining extending into axon hillocks (arrows), original magnification x 200. Western blot analysis of PrP^fl/fl^ NSECreERT transgenic mice treated with either tamoxifen (+Tx) or vehicle only (-Tx), wild type PrP expressing control sample (wt). Immunodetection of PrP protein using antibody 7A12 revealed variable reduction of brain PrP^C^ expression only after tamoxifen treatment in male (G) and female (H) mouse brain samples, GAPDH expression control from female brain samples was unchanged (I). No reduction was observed in female sciatic nerve samples (J). (K) Boxplot of brain PrP expression level following normalization to GAPDH levels and compared to wild type PrP expressing mouse. Vehicle treated mice (n=24) were statistically insignificant from wild type PrP expressing mice regardless of sex, female (n=12) and male (n=12) tamoxifen treated PrP^fl/fl^ NSECreERT mice revealed reduced PrP levels in brain, group mean indicated (♦).

### Inducible reduction of CNS PrP^C^

Having established that the anticipated recombination occurred within the CNS, we next combined the NSECreERT transgene with our gene-targeted floxed endogenous *Prnp* model (PrP^fl^), used previously in both ubiquitous (Tuzi et al., 2004) and cell-specific (Bradford et al., 2009) Cre-mediated PrP^C^ knockout models. We characterized the inducible reduction of PrP^C^ in the CNS by administering tamoxifen to PrP^fl/fl^ NSECreERT compound transgenic mice and determined the effect on total PrP^C^. There was a significant reduction in whole brain PrP^C^ post tamoxifen administration (Fig. 1G & 1H)., Different levels of PrP^C^ expression were evident between animals and between the sexes. More extensive reduction of total brain PrP^C^ was observed in the female mice (Fig. 1H & 1K) ∼45% on average, than in male mice (Fig. 1G and 1K) ∼30% on average, consistent with the beta galactosidase observations. In untreated and vehicle treated PrP^fl/fl^ NSECreERT mice, whole brain PrP^C^ levels were similar to wild type PrP expressing mice and no loss of PrP^C^ was observed (Fig. 1K), confirming that no detectable levels of sporadic Cre recombination had occurred in the absence of tamoxifen.

### Reduction of PrP^C^ in the CNS prior to infection prolongs disease incubation period

We induced PrP^C^ reduction in the PrP^fl/fl^ NSECreERT mice two weeks prior to intracerebral (i.c.) infection with the ME7 mouse-adapted scrapie (ovine TSE) strain and compared them to control groups of mice (as described in table 1). Tamoxifen treated PrP^fl/fl^ NSECreERT mice showed extended incubation periods in response to ME7 infection (Fig. 2A and Table S1). The differences in mean incubation period observed between the PrP^fl/fl^ NSECreERT groups and wild-type were statistically significant, with p values of <0.001 when compared by 2-sample t-test. PrP^C^ reduction prior to infection therefore increased disease incubation period. A greater increase was observed in female than male mice (Fig. 2A and Table S1), consistent with the greater reduction of PrP^C^ in the CNS of the female mice, indicating a reduction-response relationship. Individual assessment of the control groups revealed no statistically significant differences between males and females within these groups or between each of these groups (Fig. 2A and Table S1), indicating that individually the treatments or transgenes had no independent effect on disease incubation period.

**Table I.** Experimental groups used for the investigation of neuronal PrP^C^ depletion prior to infection.

| Group | *Prnp* | Cre | Treatment | PrP |
| --- | --- | --- | --- | --- |
| PrP depleted | fl/fl | NSECreERT | Tamoxifen | reduced |
| Vehicle only | fl/fl | NSECreERT | vehicle | normal |
| Tx/Cre -ve | fl/fl | -ve | Tamoxifen | normal |
| No treatment | fl/fl | NSECreERT | none | normal |
| 129/Ola | wt/wt | -ve | none | normal |

**Figure 2.**
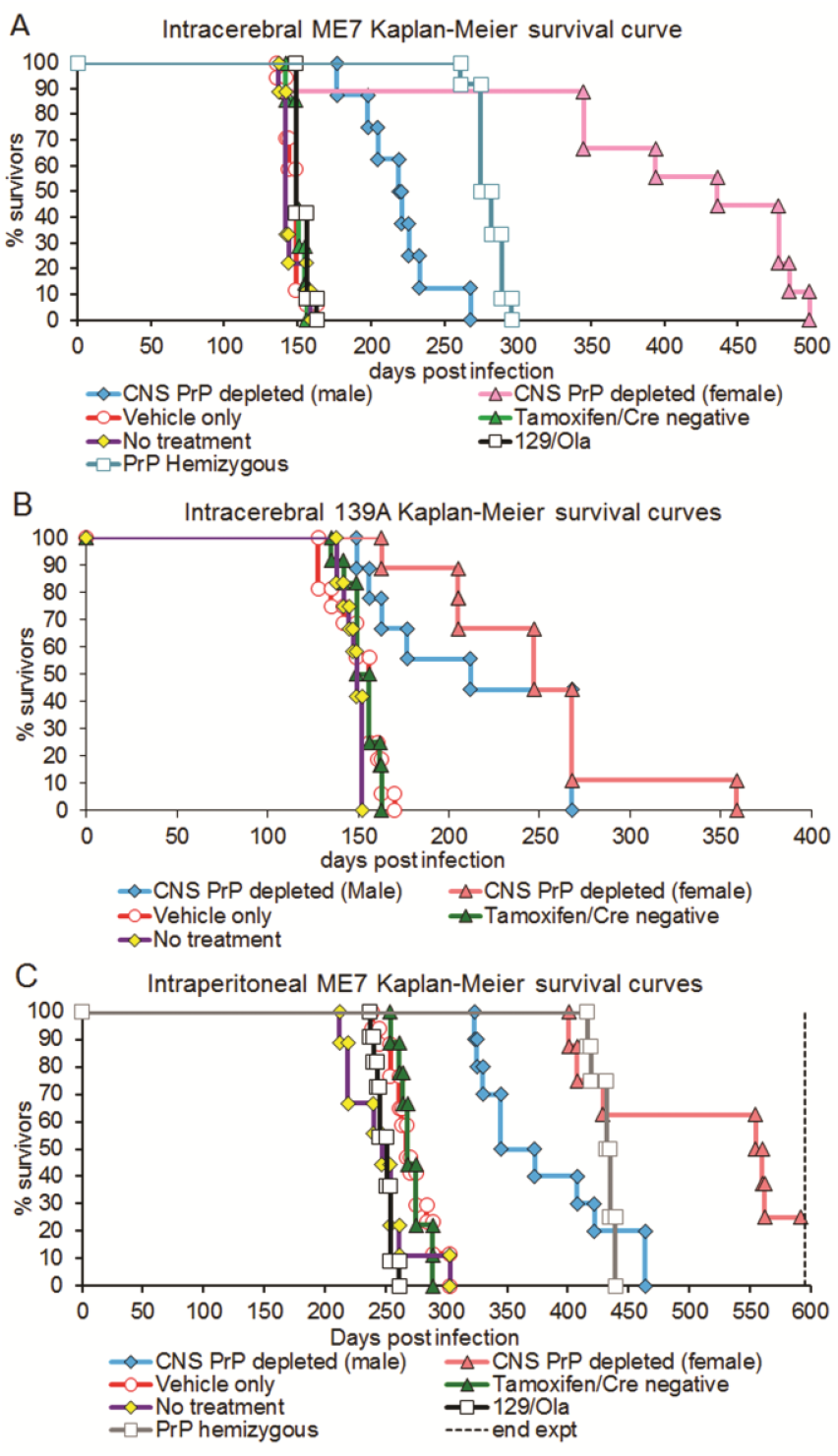
Disease incubation period analysis of PrP^C^ depletion prior to intracerebral ME7 or 139A or intraperitoneal ME7 infection. For details of each group of mice see table 1.

To establish if the effect of extended incubation period was either agent or route specific we induced PrP^C^ depletion prior to inoculation with an alternative TSE agent and using a different route of inoculation. PrP^C^ depletion was induced two weeks prior to intracerebral inoculation with mouse adapted 139A strain (Fig. 2B) and in another set of animals prior to intraperitoneal infection with the mouse-adapted ME7 strain (Fig. 2C). Tamoxifen treated PrP^fl/fl^ NSECreERT mice showed extended incubation periods in response to infection regardless of the strain of agent or route of infection (Fig. 2A-C and Table S1), although the lengthening of incubation period with 139A agent was less pronounced than that with ME7. Hence, PrP^C^ reduction increases disease incubation period irrespective of route or agent, although importantly the percentage of increase is both route and strain dependent. The differences between males and females observed following intracerebral ME7 inoculation (Fig. 2A) was also observed with intracerebral 139A strain of agent (Fig. 2B) and intraperitoneal ME7 inoculation (Fig. 2C).

### Modulation of disease incubation period throughout the preclinical phase of infection

To determine if reduction of PrP^C^ post infection could alter incubation times, we infected further cohorts of PrP^fl/fl^ NSECreERT mice, including both males and females, with intracerebral ME7 or 301C and administered the tamoxifen treatment to individual groups of mice from these cohorts at 35 and 70 days post inoculation (d.p.i.). Further groups commenced tamoxifen treatment at 105 d.p.i. (ME7) or 110 d.p.i. (301C) and the final group treated with tamoxifen coincided with the first sign of positive clinical symptoms at 126 d.p.i. (ME7 & 301C). Due to the variability observed between sexes in the group treated prior to TSE inoculation we ensured an equal sex ratio was established within each of the treatment groups. A group from each cohort was also left untreated and therefore experienced no PrP^C^ depletion; (Fig 1G & 1H). Data plotted as survival curves showing each individual mouse (Fig 3A & 3B), demonstrate a positive correlation between the proximity of induction of neuronal PrP^C^ depletion to the point of infection and the impact on mean survival period following ME7 infection. Appreciable increases in survival period (mean survival 182±15 days±SEM compared with mean survival 149±3 days±SEM of controls for ME7) were induced by PrP^C^ depletion as late as 105 days post infection, well after CNS pathologies have become established in the ME7 model (Cunningham et al., 2003, Guenther et al., 2001, Jeffrey et al., 1995). Analysis of group mean survival period revealed a statistically significant increase in ME7 incubation time from the untreated group following neuronal PrP^C^ depletion during infection in groups treated 35 and 70 days (p<0.001) post infection. In the group treated at 105 days there was a clear trend towards an increase in survival (p=0.062). Following 301C infection an even greater effect was observed with induction of PrP^C^ depletion at any time prior to clinical symptoms (35, 70 and 110 d.p.i.) revealing similar increases in mean survival time (300±30, 264±29 and 279±36 days±SEM respectively compared to mean survival of non-depleted controls 153±4 days±SEM). The extended incubation times in the preclinical phase of 301C appeared less dependent on the stage of disease with greatly extended time occurring throughout the preclinical phase. Where PrP^C^ reduction was concurrent with the onset of clinical symptoms of TSE disease, similar incubation periods to control non-tamoxifen treated mice expressing PrP^C^ at the same levels as wild type were found regardless of the strain of agent, demonstrating that at this stage of infection the incubation period of disease was not affected by the reduced levels of PrP^C^.

**Figure 3.**
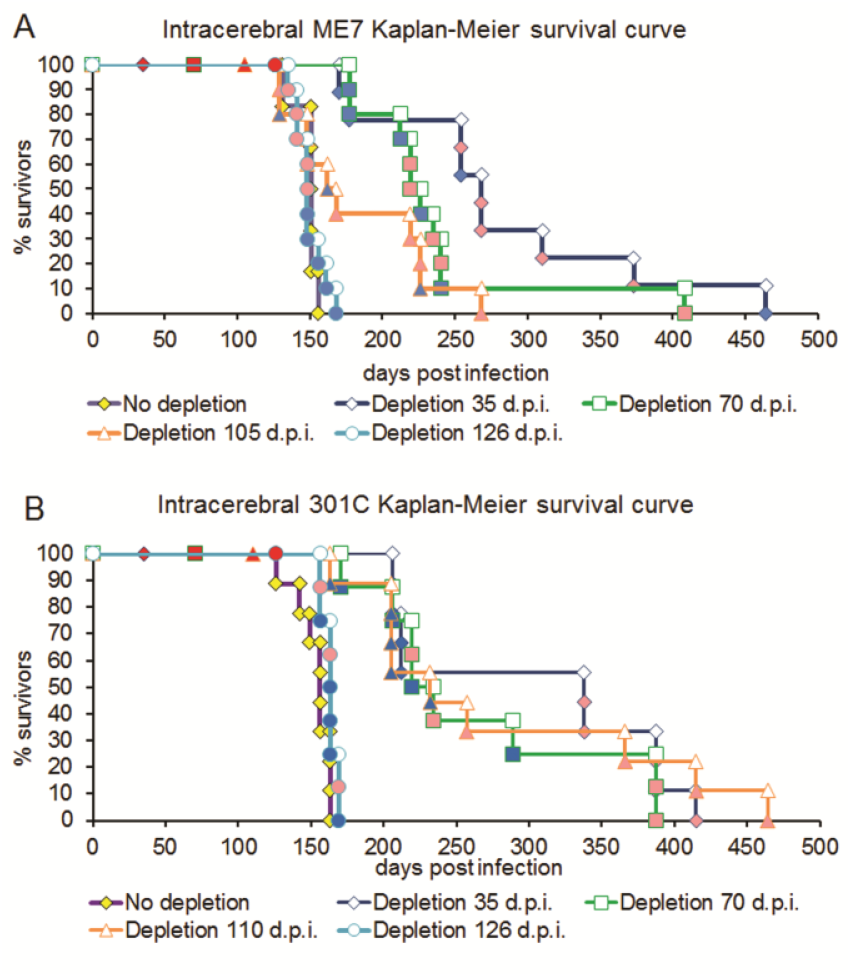
Disease incubation period analysis of PrP^C^ depletion during intracerebral ME7 or 301C infection. (A) Intracerebral ME7 and (B) intracerebral 301C Kaplan-Meier survival curves with PrP^C^ depletion during infection. Red symbols indicate the timing of PrP^C^ depletion, pink symbols represent female mice and blue symbols male mice in each data series.

### CNS PrP^C^ depletion alters TSE pathogenesis and promotes neuroprotection

We examined the pathological lesions in mice treated with neuronal PrP^C^ depletion prior to infection and compared the ME7 infected wild type PrP mice with the PrP^fl/fl^ NSECreERT mice. Brain sections from terminal ME7-infected PrP^fl/fl^ NSECreERT animals showed a striking preservation of neuronal soma in the CA1 region of the dorsal hippocampus (Fig. 4I & 4M), an area which displays heavy neuronal loss following ME7 infection in wild type PrP expressing mice (Fig. 4A). Individuals with the longest disease incubation time had a neuronal density within the CA1 equivalent to control mice, despite displaying clinical symptoms and succumbing to TSE disease. A reduction in the thickness of the hippocampal layers was observed in terminal ME7 infected wild type PrP expressing mice (Fig. 4A). However, terminal ME7-inoculated PrP^fl/fl^ NSECreERT mice revealed a preservation of the hippocampal layer thickness (e.g. stratum radiatum) (Fig. 4I & 4M).

**Figure 4.**
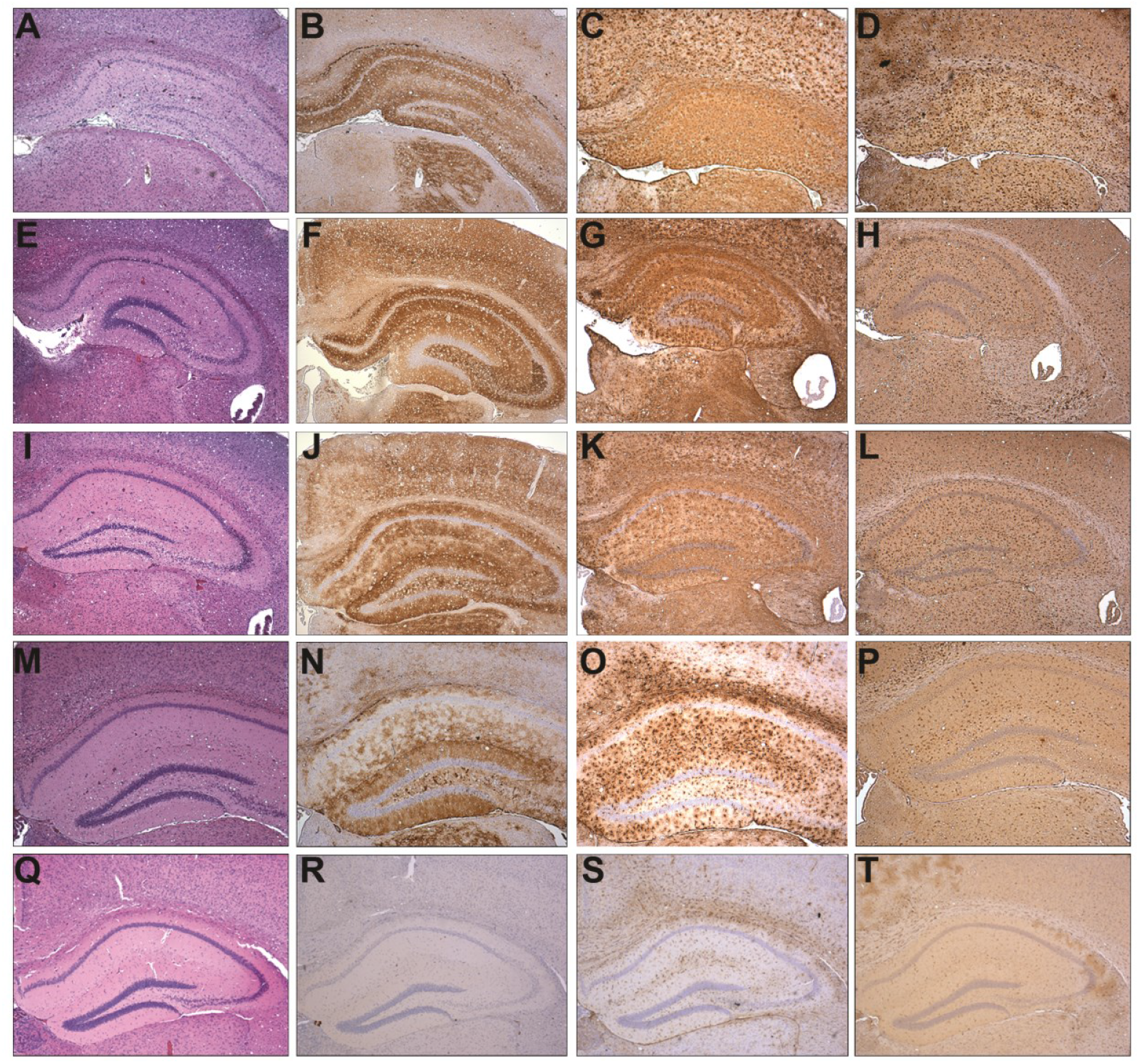
Depletion of PrP^C^ from neurones prior to intracerebral ME7 infection alters TSE-induced hippocampal neuropathology. Comparison of terminal ME7 neuropathology in PrP wild type (A-D), hemizygous PrP null (E-H) and tamoxifen treated PrP^fl/fl^ NSECreERT male (I-L) and female (M-P) mice reveals alteration to common TSE neuropathology not observed in uninfected wild type mice (Q-T). (A, E, I, M & Q) haematoxylin and eosin stained sections; (B, F, J, N & R) anti-PrP antibody 6H4; (C, G, K, O & S) anti-GFAP staining of astrocytes and (D, H, L, P & T) anti-AIF1 staining of microglia. Terminal ME7 infected PrP wild type mice revealed extensive loss of neurones (A), widespread PrP^Sc^ deposition (B) and reactive astro-(C) and micro-gliosis (D). Hemizygous PrP null mice reveal widespread vacuolation with reduced neuronal loss (E), PrP^Sc^ deposition (F) and reactive gliosis (G & H). Tamoxifen treated PrP^fl/fl^ NSECreERT of both sexes revealed altered targeting of vacuolation predominantly to the dentate gyrus within the hippocampus and marked preservation of pyramidal neurone soma (I & M), despite extensive in male (J), or patchy PrP^Sc^ deposition in females (N). Glial responses to infection were moderately altered in males (K & L) or markedly altered in females (O & P). Original magnification of all images = x40.

Terminal ME7 infected wild type PrP expressing mice displayed widespread deposition of disease-associated PrP^Sc^ in particular in the hippocampus and cerebral cortex layer IV. Intense PrP deposition was observed in the form of fine-punctate and coarse deposits throughout the stratum radiatum and lacunosum moleculare with plaque-like deposits in the corpus callosum (Fig. 4B). Despite the reduction of PrP^C^, extensive deposition of PrP was observed in the brains of the terminal ME7 infected tamoxifen treated PrP^fl/fl^ NSECreERT mice (Fig. 4J & 4N). The pattern in groups with the most extensive reduction of PrP^C^ was dramatically different to that of the wild type mice PrP deposition, appeared to associate with astrocytes rather than neurons, especially in the neuropil strata and pyramidal layers of the hippocampus, cortex and thalamus while deposition in the dentate gryus of these animals was diffuse (Fig. 4N). Following intraperitoneal challenge similar alterations to pathology were observed in response to PrP^C^ depletion. To investigate this altered pattern of deposition of PrP in the CNS a more extensive pathological investigation was conducted in mice with the greatest reduction in PrP i.e. the female mice.

### Neuronal PrP^C^ depletion preserves hippocampal dendro-synaptic markers but reveals structural disorganization

Investigations of the dendritic arborization and synaptic cytoarchitecture of the hippocampal CA1 regions were carried out via immunostaining for the neuronal specific microtubule associated protein isoforms 2A+2B (MAP2; uniprot P20357, Fig. 5A, 5C & 5D) and for the pre-synaptic vesicle protein synaptophysin (SYP; uniprot Q62277, Fig 5B, 5E & 5F). Terminal ME7 infected female tamoxifen treated PrP^fl/fl^ NSECreERT mice revealed a high degree of preservation of the quantity of both markers within CA1 region (Fig. 5D & 5F), with the staining intensity observed similar to uninfected mice (Fig. 5A & 5B). A moderate degree of disorganization was apparent for both markers suggesting that some disruption has occurred to the dendritic architecture of the neurons in this region (Fig. 5D & 5F) compared to the fine dendritic projections observed in uninfected mice (Fig. 5A and 5B). Terminal ME7-infected wild type PrP^C^ expressing mice revealed extensive disruption of the distribution or loss of MAP2 (Fig. 5C) and SYP (Fig. 5E) concurrent with the loss of neuronal soma (Fig. 4A) when compared to uninfected mice (Fig. 5A & 5B).

**Figure 5.**
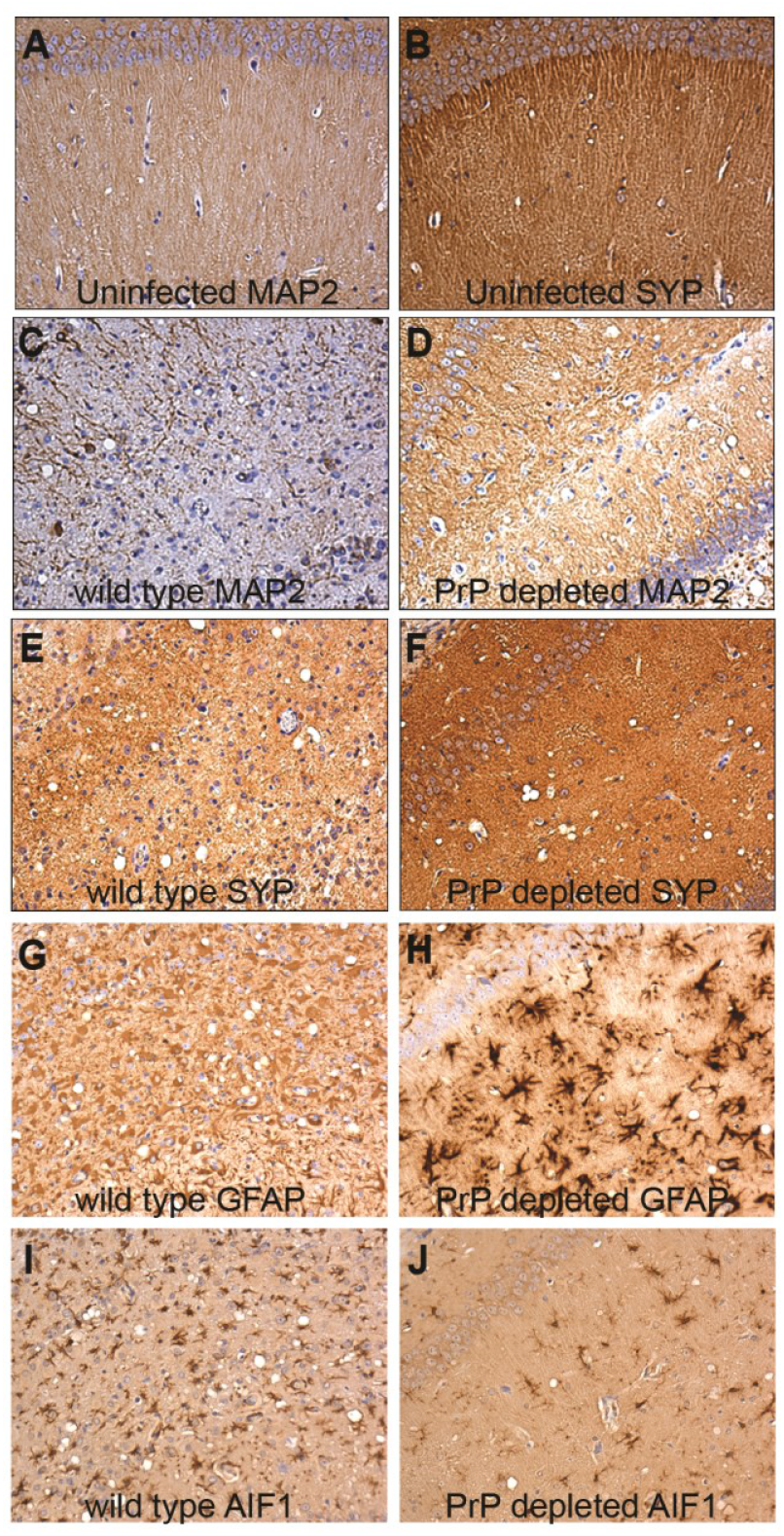
Depletion of PrP^C^ from neurons prior to intracerebral ME7 infection alters CA1 neuropathology. Comparison of terminal ME7 neuropathology in PrP wild type (C, E, G & I) and PrP^fl/fl^ NSECreERT mice (D, F, H & J) compared to uninfected mice (A & B). Typical CA1 dendritic arborization and synapse density in uninfected mice is revealed via immunohistochemical staining for MAP2 (A) and SYP (B). Terminal ME7 infected PrP wild type mice revealed extensive loss of dendritic (C) and synaptic (E) markers from CA1 pyramidal neurones. Within the hippocampal CA1 region, terminal ME7 infected female tamoxifen treated PrP^fl/fl^ NSECreERT mice reveal retention of neuronal dendritic (D) and pre-synaptic markers (F) but with some disorganization. Following ME7 infection GFAP+ astrocytes in the CA1 region of female tamoxifen treated PrP^fl/fl^ NSECreERT mice appear fibrous (H), unlike the gemistocytic appearance of CA1 astrocytes in wild type PrP expressing mice (G). In contrast, fewer morphologically reactive (Aif1+) microglia were observed within the layer of pyramidal neuron soma of female tamoxifen treated PrP^fl/fl^ NSECreERT mice (J), unlike in PrP wild type mice (I) at the terminal stage of ME7 infection. Original magnification of all images = x200.

### Neuronal PrP^C^ depletion alters glial response to TSE infection

In terminal ME7 infected wild type mice GFAP immunostaining produced a heavy but diffuse pattern throughout the hippocampal layers; with hypertrophied astrocytes in heavily affected disease areas and distinct gemistocytic appearance particularly within the stratum radiatum and lacunosum moleculare (Fig. 4C & 5G). In contrast in terminal ME7 infected female tamoxifen treated PrP^fl/fl^ NSECreERT mice fibrous, hypertrophic astrocytes were observed in all hippocampal regions following GFAP immunostaining (Fig. 4O & 5H). Immunostaining for AIF1 revealed diffuse microgliosis in all layers of the hippocampus in wild type mice at terminal stage of ME7 infection (Fig. 4D & 5I). Reactive AIF1^+^ microglia were also observed throughout the hippocampus of terminal ME7 infected female tamoxifen treated PrP^fl/fl^ NSECreERT mice, but few were observed within the pyramidal cell layer of the hippocampus and the density of reactive microglia was greater in the dentate gyrus than the stratum radiatum and lacunosum moleculare (Fig. 4P & 5J).

## Discussion

In the PrP^fl/fl^ NSECreERT mice, reduction of PrP in the CNS prior to TSE infection led to extensive alterations to incubation period, groups with the lowest levels of PrP^C^ expression within the CNS being associated with the longest incubation times. Depletion of neuronal PrP throughout the subclinical phase of disease resulted in prolonged incubation times. Pathogenesis was delayed and pathological lesions altered, but disease was not arrested and clinical symptoms and death ultimately occurred. Modulation of the disease process by manipulating protein expression would appear to be possible during the preclinical period, even when substantial pathological signs of disease are present (Cunningham et al., 2003, Guenther et al., 2001, Jeffrey et al., 1995), providing a wide window for therapeutic intervention.

The lowest observed level of PrP^C^ in tamoxifen treated PrP^fl/fl^ NSECreERT mice was a 70% reduction (Fig. 1), representing a substantial knockdown of the PrP^C^. Residual PrP in the CNS could arise from two possibilities, from an incomplete removal of neuronal PrP or from expression of PrP in other cell types. The altered pathological lesions observed in this study (Fig. 4 & 5) suggest that the production of PrP^C^ from non-neuronal cells should be considered as an important contributor to the disease process. To establish possible non-neuronal sources of the residual PrP in these mice, we profiled the levels of *Prnp* mRNA in both neuronal and non-neuronal cells, using publicly available results from microarray analysis (Supplementary material). These analyses show that *Prnp* is expressed extensively in neuronal tissues and cell types, but the highest normalised expression is in astrocytes of mice (postnatal days 17 and 30) (Fig. S1). Thus, astrocytes appear to express substantial amounts of *Prnp* mRNA and therefore have the potential to contribute significant levels of PrP protein in the brain and also to be actively involved in the disease process through their production of PrP. Global removal of PrP results in animals that are resistant to infection (9-11), while in our animals removal of PrP in neuronal cells led to prolonged incubation times but not to absence of disease, suggesting that astrocytic production of PrP would have to be taken into account in any therapeutic strategy.

Neuronal cell bodies in the hippocampus, normally a major target for degeneration in the ME7 model, were protected when PrP was depleted from neurons. Although the neuronal cell bodies appeared intact we observed disruption of dendritic processes in the stratum radiatum which may result from extensive astrocytic associated build-up of protein or perhaps from astrocytic associated neurodegenerative mechanisms targeting synapses, dendrites and axons rather than cell bodies. The dramatically altered pathology in the hippocampus of the female tamoxifen treated PrP^fl/fl^ NSECreERT mice when compared with wild type mice is evidence that depletion of neuronal PrP has profound effects on the disease process. The microglial responses in this region of PrP depleted mice were much reduced compared to wild type mice, suggesting a lack of activation of microglia despite the astrocytic response and build-up of misfolded protein.

In conclusion, PrP reduction throughout the preclinical phase of disease can dramatically extend the incubation time but does not prevent the ultimate development of a neurological disease. Importantly, this study has provided evidence for a wide therapeutic window for disease intervention. Understanding the interaction of non-neuronal and neuronal cells during the course of disease is now a key factor in defining the mechanisms of prion diseases and other protein misfolding disease such as Alzheimer’s Disease, Parkinson’s disease and amyotrophic lateral sclerosis (Ilieva et al., 2009) but such a wide therapeutic window provides significant encouragement that treatment should be possible. Our study has important implications both for prion diseases induced by an infectious agent or genetic factors and for other protein misfolding conditions such as Alzheimer’s disease induced by genetic or environmental factors, since it indicates that early identification of at risk individuals and prompt treatment (prior to or even during the subclinical phase) could delay the disease process potentially well beyond the lifetime of an individual.

## Materials and Methods

### Generation of inducible neuronal Cre mice

A 4kb fragment containing the *Rattus norvegicus* neuron specific gamma enolase (*Eno2 gene;* uniprot P07323) promoter (AB038993) was excised from NSE-n1.HPrP plasmid (Race et al., 1995) using HindIII and SalI, gel purified and sub-cloned into pBluesript propagated in XL-1 Blue competent cells. A further fragment was excised with MfeI and NheI, gel purified and ligated into the PCre5orr plasmid upstream of the Cre modified estrogen receptor fusion protein sequence (CreERT) pre-cut with the same enzymes, and transfected into XL-1 Blue cells. A 6.9kb fragment containing the whole NSECreERT minigene was excised with MfeI and SphI and microinjected into the pronucleus of C57Bl/6 fertilized embryos. Founder mice were screened via PCR for the presence of the CreERT fusion protein sequence with the oligonucleotide primers 5’-ATACCGGAGATCATGCAAGC-3’ and 5’-CAAAGCCTGGCACTCTCTTT-3’.

### Generation of inducible neuronal PrP depleted transgenic mice

Mice positive for the NSECreERT transgene were selectively backcrossed against Prnp^tm2Tuzi^ (PrP^fl^) mice (Tuzi et al., 2004). Offspring devoid of the NSECreERT transgene but homozygous for PrP^fl^ were also identified and inbred to create a Cre negative control line with similar genetic background (see table 1).

### Tamoxifen administration

Tamoxifen (Sigma) was dissolved in sunflower seed oil pre-heated to 80°C at a concentration of 10 mg ml^-1^. Mice were dosed by bodyweight for five consecutive days at 0.08 mg g^-1^. As controls, groups of mice of the same genotype received an equivalent volume of sunflower seed oil vehicle only, no treatment at all, or tamoxifen dosage with a PrP^fl/fl^ genotype lacking the NSECreERT transgene (see table 1).

### Transmissible spongiform encephalopathy infection and clinical observation

Pooled brain tissue from terminally ME7, 139A or 301C infected mice were homogenized at 10 % w/v in saline solution and stored frozen at -40°C until use. Inoculum was diluted to 10^−2^ with saline solution upon thawing. Animals were inoculated with 20 µl inoculum intracerebrally (i.c.) or 100 µl intraperitoneally (i.p.). Intracerebral inoculations were performed while mice were under isofluorane-induced light anesthesia. Mice were clinically observed weekly as described previously (Bruce et al., 1991) starting from 100 days post-inoculation. Disease incubation times were calculated as the interval between inoculation and positive clinical assessment of terminal TSE disease and post mortem confirmation by positive neuropathological assessment.

### Ethics Statement

All animal studies and breeding were performed via standard methods and conducted under the provisions of the UK Animals Scientific Procedures Act 1986 and approved by the Neuropathogenesis Division Ethical Review Committee. Animals were housed under standard conditions with 12 hour light/dark cycle and food and water ad libitum. Animals were monitored daily for signs of ill-health or distress and clinically observed weekly as described previously (Bruce et al., 1991) starting from 100 days post TSE inoculation. Animals revealing signs of ill-health other than clinical symptoms of TSE disease were euthanized immediately for ethical reasons and full pathological assessments were performed, including neuropathology. All animals were coded to blind the experimenter to genotype, treatment and inoculum groups.

### Western immunoblot analysis

Western immunoblot analysis was performed as described previously (Bradford et al., 2009), briefly Ten per-cent (w/v) brain and sciatic nerve homogenates were prepared and immunoblotted with primary antibodies at dilutions indicated: monoclonal Mouse anti PrP clone 7A12 1:20,000; monoclonal Mouse Anti-Glyceraldehyde-3-Phosphate Dehydrogenase (GAPDH) clone 6C5 (Millipore) 1:10,000.

### Neuropathological assessment

Tissue for neuropathological assessment was coded to allow blind-scoring and fixed in 10% formal saline for a minimum of 48 hr, processed and embedded in paraffin. Hematoxylin and eosin stained 6 µm sections were assessed and scored for spongiform vacuolar degeneration on a scale of 0 to 5 grey matter or 0 to 3 white matter in 12 standard areas as previously described (Fraser and Dickinson, 1967).

### Histochemistry and immunohistochemistry

X-gal histochemistry and immunohistochemistry were performed as described previously (Bradford et al., 2009). Briefly sections were incubated with primary antibody overnight at dilutions indicated: monoclonal Mouse anti-PrP clone 6H4 (Prionics) 1:1,000, polyclonal Rabbit anti-GFAP (Dako) 1:1,000; polyclonal Rabbit anti-iba1 (Wako) 1:1,000; monoclonal Mouse anti-Synaptophysin clone SY38 (DAKO) 1:500; monoclonal Mouse anti Microtubule associated proteins 2A+2B (Abcam) 1:1,000. Sections were imaged on an E800 microscope (Nikon) using Image pro plus software (Media Cybernetics).

### Statistical analysis

Group mean incubation periods were calculated and compared by analysis of variance (ANOVA) for each experiment. Kaplan-Meier curves were constructed censored on animals displaying both positive clinical and pathological assessments. 2 sample t-tests were performed for groups with extended incubation period versus combined control groups expressing PrP normally. Group mean vacuolation scores were plotted against area to give representative profiles. Neuronal counts were performed using ImageJ software. Statistical analyses were performed using Minitab software and all graphs plotted using Excel (Microsoft).

## Acknowledgements

We thank colleagues at the Neuropathogenesis Division of the Roslin Institute and R(D)SVS the University of Edinburgh, in particular Irene McConnell, Simon Cumming, Kris Hogan and Val Thomson for the breeding, care and maintenance of transgenic mouse lines and the setup and scoring of TSE challenge experiments; Anne Coghill, Aileen Boyle, Sandra Mack and Gillian McGregor for histological processing and vacuolation scoring. NSE-n1.HPrP plasmid kindly provided by Bruce Chesebro (Rocky Mountain Laboratories, Montana, USA). PCre5orr plasmid kindly provided by Dr Tom Gardner (University of Edinburgh, UK). Antibody 7A12 was kindly provided by Man-Sun Sy (Case Western Reserve University, Ohio, USA). ROSA26-LacZ mice were kindly provided by Drs Yuko Fujiwara and Stuart H. Orkin (Harvard Medical School, Massachusetts, USA).

## Author Contributions

Conceived and designed the experiments: BMB AM NLT KMS VHP ARC JCM. Performed the experiments: BMB AM DB DK KMS. Analyzed the data: BMB DB KMS PP. Wrote the paper: BMB KMS PP JCM.

