## Supplementary material for "Therapeutic window for treatment of prion disease defined by timed depletion of Prion protein": Supplemetary data

**Supplementary Table 1: Response to infection; group mean incubation period.**

| Group | Inocula | Route | Sex | N= | Incubation period (Days +/- SEM) | Survival |
|---|---|---|---|---|---|---|
| PrP depleted | ME7 | i.c. | Male | 6 | 229±9 | |
| | | | Female | 10 | 331±32 | |
| Vehicle only | ME7 | i.c. | Male | 10 | 149±2 | |
| | | | Female | 7 | 144±2 | |
| Tx/Cre -ve | ME7 | i.c. | Male | 2 | 153±2 | |
| | | | Female | 5 | 149±2 | |
| No treatment | ME7 | i.c. | Male | 3 | 141±2 | |
| | | | Female | 6 | 147±3 | |
| 129/Ola | ME7 | i.c. | Male | 8 | 153±2 | |
| | | | Female | 4 | 153±2 | |
| C57Bl/6 | ME7 | i.c. | Male | 2 | 156±0 | |
| | | | Female | 4 | 156±0 | |
| PrP hemizygous | ME7 | i.c. | Male | 8 | 283±3 | |
| | | | Female | 4 | 275±6 | |
| PrP depleted | ME7 | i.p. | Male | 10 | 386±17 | |
| | | | Female | 8 | 467±30 | 2 @ 609 |
| Vehicle only | ME7 | i.p. | Male | 10 | 261±5 | |
| | | | Female | 7 | 283±6 | |
| Tx/Cre -ve | ME7 | i.p. | Male | 4 | 266±4 | |
| | | | Female | 5 | 288±16 | |
| No treatment | ME7 | i.p. | Male | 4 | 237±10 | |
| | | | Female | 5 | 253±15 | |
| 129/Ola | ME7 | i.p. | Male | 7 | 250±3 | |
| | | | Female | 4 | 246±3 | |
| PrP hemizygous | ME7 | i.p. | Male | 5 | 433±4 | |
| | | | Female | 3 | 428±6 | |
| PrP depleted | 139A | i.c. | Male | 9 | 214±18 | |
| | | | Female | 9 | 248±18 | |
| Vehicle only | 139A | i.c. | Male | 11 | 157±5 | |
| | | | Female | 5 | 157±1 | |
| No treatment | 139A | i.c. | Male | 6 | 149±2 | |
| | | | Female | 6 | 145±3 | |
| Tx/Cre -ve | 139A | i.c. | Male | 9 | 154±2 | |
| | | | Female | 3 | 149±8 | |
| PrP depleted | none | n/a | Male | 11 | | 411±51 |
| | | | Female | 12 | | 551±55 |

**Supplementary Table 2: Datasets used to examine gene expression profiles.**

| GEO DataSets Accession Number | Cell type | Reference |
|---|---|---|
| GSM258651, GSM258652 | Dorsal root ganglia | (Lattin et al., 2008) |
| GSM258635, GSM258636 | Cerebral cortex | (Lattin et al., 2008) |
| GSM358637, GSM258638 | Prefrontal cortex | (Lattin et al., 2008) |
| GSM258735, GSM258736 | Olfactory bulb | (Lattin et al., 2008) |
| GSM258733, GSM258734 | Nucleus accumbens | (Lattin et al., 2008) |
| GSM258653, GSM258654 | Dorsal striatum | (Lattin et al., 2008) |
| GSM258617, GSM258618 | Amygdala | (Lattin et al., 2008) |
| GSM258671, GSM258672 | Hippocampus | (Lattin et al., 2008) |
| GSM258633, GSM258634 | Cerebellum | (Lattin et al., 2008) |
| GSM258673, GSM258674 | Hypothalamus | (Lattin et al., 2008) |
| GSM258749. GSM258750 | Pituitary | (Lattin et al., 2008) |
| GSM258757, GSM258758 | Retina | (Lattin et al., 2008) |
| GSM258765, GSM258766 | Spinal cord | (Lattin et al., 2008) |
| GSM241896, GSM241904 | Neurons P16 | (Appolloni et al., 2009) |
| GSM241908 | Neurons P26 | (Appolloni et al., 2009) |
| GSM258727, GSM258728 | Neuro2a cell line | (Lattin et al., 2008) |
| GSM387042, GSM387043, GSM387044, GSM387045 | Oligodendrocytes | (Emery et al., 2009) |
| GSM241912, GSM241914, GSM241926 | Astrocytes P17 | (Appolloni et al., 2009) |
| GSM241919 | Astrocytes P30 | (Appolloni et al., 2009) |
| GSM258721,GSM258722 | Microglia | (Lattin et al., 2008) |
| GSM258693, GSM258694 | Bone marrow macrophages | (Lattin et al., 2008) |

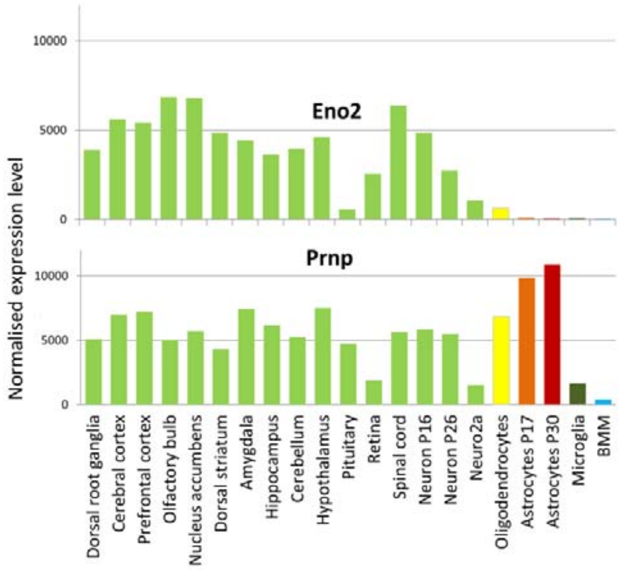

**Figure S1. Gene expression analysis of Prnp and Eno2 in CNS cell types**

Publicly available gene expression data generated from mouse neuronal tissues and cell types using the Affymetrix MOE430_2 GeneChip were accessed through GEO DataSets (Supplementary material). Downloaded CEL files were normalized by the robust microarray averaging (RMA) method using the Affymetrix Gene Expression Console. Normalized untransformed results were used in the analysis. Y axis shows the normalized gene expression levels for the different cells and tissues shown on the X axis. Data reveals that highest *Prnp* expression levels were observed in astrocytes as opposed to individual neuronal populations throughout the CNS, and that *Eno2* expression levels vary between different brain areas.
